# Fine-tuning of the AMBER RNA Force Field with a New Term Adjusting Interactions of Terminal Nucleotides

**DOI:** 10.1101/2020.03.08.982538

**Authors:** Vojtěch Mlýnský, Petra Kührová, Tomáš Kühr, Michal Otyepka, Giovanni Bussi, Pavel Banáš, Jiří Šponer

## Abstract

Determination of RNA structural-dynamic properties is challenging for experimental methods. Thus atomistic molecular dynamics (MD) simulations represent a helpful technique complementary to experiments. However, contemporary MD methods still suffer from limitations of force fields (*ff*s), including imbalances in the non-bonded *ff* terms. We have recently demonstrated that some improvement of state-of-the-art AMBER RNA *ff* can be achieved by adding a new term for H-bonding called gHBfix, which increases tuning flexibility and reduces the risk of side-effects. Still, the first gHBfix version did not fully correct simulations of short RNA tetranucleotides (TNs). TNs are key benchmark systems due to availability of unique NMR data, although giving too much weight on improving TN simulations can easily lead to over-fitting to A-form RNA. Here we combine the gHBfix version with another term called tHBfix, which separately treats H-bond interactions formed by terminal nucleotides. This allows to refine simulations of RNA TNs without affecting simulations of other RNAs. The approach is in line with adopted strategy of current RNA *ff*s, where the terminal nucleotides possess different parameters for the terminal atoms than the internal nucleotides. The combination of gHBfix with tHBfix significantly improves the behavior of RNA TNs during well-converged enhanced-sampling simulations. TNs mostly populate canonical A-form like states while spurious intercalated structures are largely suppressed. Still, simulations of r(AAAA) and r(UUUU) TNs show some residual discrepancies with the primary NMR data which suggests that future tuning of some other *ff* terms might be useful.

## INTRODUCTION

Atomistic description of structural dynamics of RNA molecules is essential for understanding of some of key biomolecular processes including, e.g., proteosynthesis, splicing, ribosomal catalysis etc. Experimental techniques, such as X-ray crystallography, cryo-electron microscopy and nuclear magnetic resonance (NMR) spectroscopy provide essential structural information. However, the dynamics is obscured by ensemble averaging. Insights into biomolecular motions and structural dynamics can also be obtained by molecular dynamics (MD) simulations. However, currently used (state-of-the-art) empirical potentials (force fields, *ff*s) suffer from many inaccuracies which may even lead to a preferential sampling of spurious structures instead of the native states. Performance of the available pair-additive atomistic RNA *ff*s has been reviewed in detail,^1–4^ with a suggestion that the currently used form of the pair-additive RNA *ff*s is approaching the limits of its applicability.^4^ In other words, it is becoming increasingly difficult to generally improve performance of the RNA simulations by refining parameters of the available *ff* terms without introducing some new unintended imbalances. Radical approach to overcome the limitations is the ongoing development of more sophisticated polarizable *ff*s.^5–9^ Alternatively, we have suggested that improvements could be achieved also by extending the pair-additive *ff*s by new simple terms that are uncoupled from the existing *ff* terms and thus allow tuning of simulations of problematic systems while minimizing undesirable side effects. We have introduced an additional *ff* term called HBfix (H-bond fix),^10^ that can be used to tune the description of key hydrogen-bonding interactions. HBfix is a simple short-range potential (typically acting in the range from 2 to 3 Å for H – acceptor distances) that either increases or decreases stability of target H-bonds (Figure 1). It has been first applied as a structure-specific potential which can improve folding of some RNA tetraloops^10^ and structural stability of protein-RNA complexes.^11^ Subsequently, we have suggested its generalized interaction-specific variant (gHBfix) which represents a true extension of the parent *ff*.^12^ We have demonstrated that boosting of all –NH…N– base–base interactions by 1.0 kcal/mol and destabilizing all sugar–phosphate (SPh) H-bonds by 0.5 kcal/mol significantly improves simulations of RNA tetranucleotides and GNRA tetraloop without deteriorating simulations of other systems (Figure 1). These parameters complement the common OL3 (*ff*99bsc0χ_OL3_)^13–16^ AMBER RNA *ff* version used together with modified phosphate van der Waals (vdW) parameters^17^ and OPC water model.^18^

**Figure 1:**
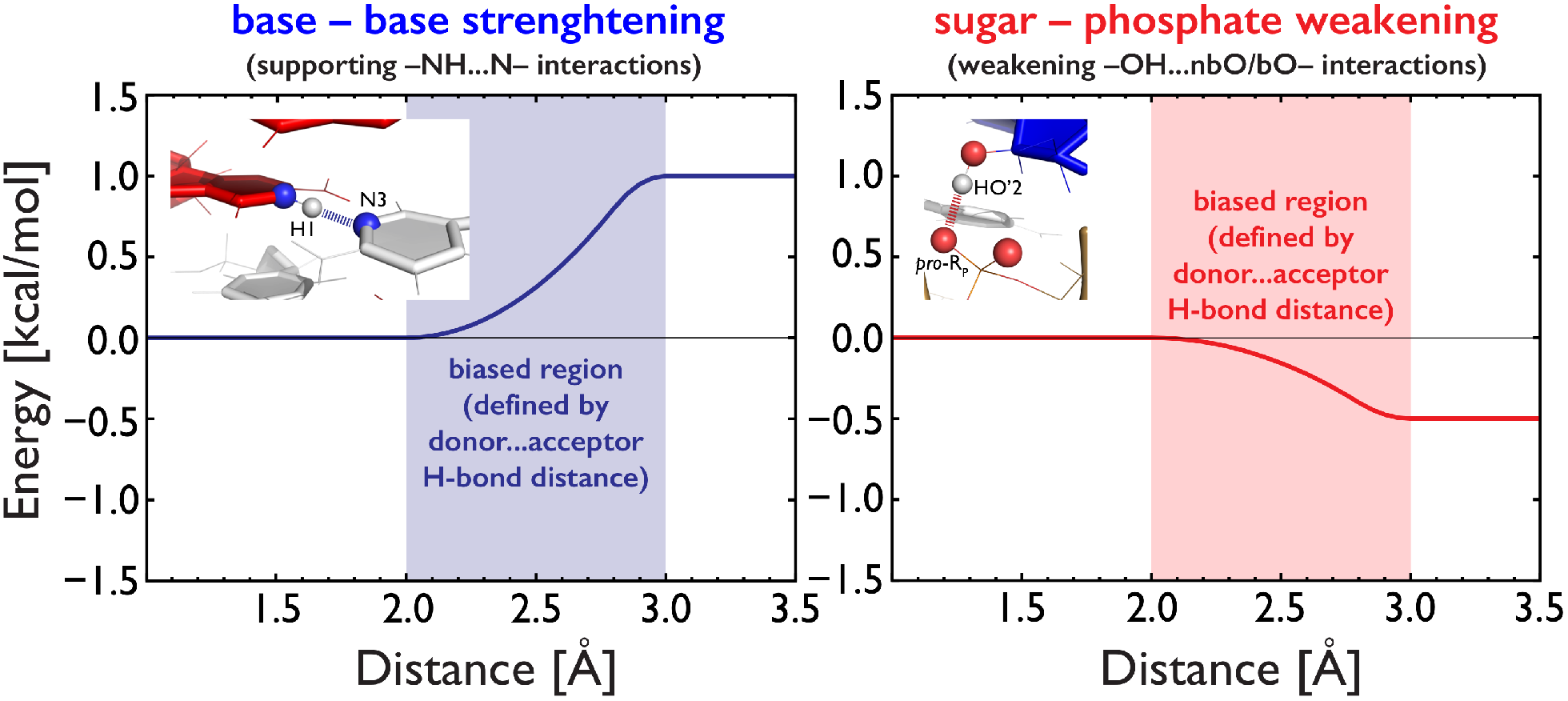
Description of the gHBfix *ff* term used for either support (blue curve) or weakening (red curve) of H-bond interactions. The initial version of the gHBfix introduced in Ref. ^12^ is tuning H-bond interactions in order to: (i) support base – base interactions (left panel, blue dashed lines for –NH…N–interactions) and, simultaneously, (ii) destabilize SPh interactions (red dashed lines on the right panel); SPh interactions are those between 2’-OH groups and bridging (bO)/non-bridging (nbO) phosphate oxygens. The gHBfix potential is affecting all interactions of the same type while an analogous term can be used to modulate only structure-specific (individual) H-bonds (abbreviated as HBfix^10, 12^). In the present work, we suggest a variant separately targeting interactions involving terminal nucleotides (tHBfix term).

In this work we show that further improvement of RNA simulations can be achieved by tuning the SPh and base-phosphate (BPh) H-bonds involving the terminal nucleotides; this variant is henceforth abbreviated as tHBfix. This approach, although targeting only selected nucleotides can be considered as a generally transferable *ff* modification, as it can be applied to all simulated RNAs. Tuning of parameters of terminal nucleotides in RNA chains is fully justified as their properties can differ from internal nucleotides and their imbalanced description in simulations can have detrimental effects. In other words, modification of the interactions of terminal nucleotides can separately eliminate some simulation problems without affecting the remaining parts of the RNA molecules. This can reduce the risk of overfitting the *ff* when using simulations of small model RNA systems for its testing and training. The rationale is that terminal nucleotides have excessively large impact on simulations of small model systems and thus it is advisable to screen off their role as much as possible. Our ultimate goal is not to reproduce experimental data for these small model systems, but to study folded RNA structures and diverse protein-RNA complexes. Targeting the terminal residues of the *ff* may be considered both as an attempt to reflect potentially different electronic structures of these residues and a pragmatic means to increase tuning capability of the *ff*.

The present study is based mainly on simulations of RNA tetranucleotides (TNs). Conformation ensembles of TNs provide one of the key benchmarks for testing RNA *ff*s due to their small size and straightforward comparison of their simulations with solution experiments.^19–31^ Obviously, any quantitative *ff* assessment is critically dependent on the convergence of structural populations because only well-converged simulations can provide unambiguous benchmark datasets. Nevertheless, contemporary simulation methods and hardware already allow to obtain sufficiently converged simulation ensembles for TNs.^12, 20, 30^ TN simulations can specifically monitor performance of *ff*s for several salient energy contributions, namely, (i) SPh and BPh interactions, (ii) base stacking interactions, (iii) backbone conformations and (iv) balance of these contributions with solvation. Experimental data shows that TNs mostly populate A-form conformations while MD simulations tend to significantly sample also non-native intercalated structures (or some other non-native structures, Figure 2) that are considered to be *ff* artifacts.^12, 20, 24, 30^ Obviously, when using TNs as a benchmark for *ff* refinement, one has to be concerned about a possible over-fitting of the *ff* towards the canonical A-RNA conformation (a potential problem of some recently published *ffs*^32^), which may lead to side-effects for simulations of folded RNAs.^12, 33^

**Figure 2:**
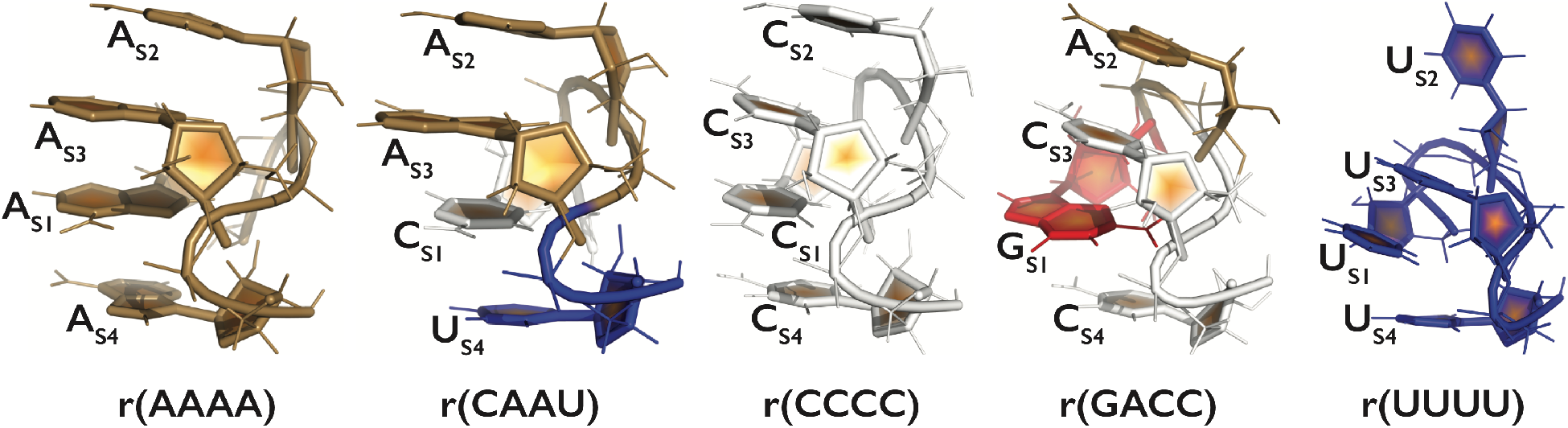
Tertiary structures of the five studied systems in the spurious intercalated structure involving the 5’-terminal nucleotide, which is often seen during MD simulations using common RNA AMBER *ff*s.^12, 19–25, 27, 28, 30–32^ A, C, G and U nucleotides are colored in sand, white, red, and blue, respectively.

Here, we applied enhanced-sampling simulations to obtain conformational ensembles of five RNA tetranucleotides, i.e., r(AAAA), r(CAAU), r(CCCC), r(GACC), and r(UUUU). The basic gHBfix potential combined with additional fixes involving one or both terminal residues (tHBfix, Table 1) significantly increases agreement between predicted and experimental data. The obtained conformational ensembles revealed that r(CAAU), r(CCCC) and r(GACC) TNs are sampling more than 75% of time A-form–like conformations. The remaining r(AAAA) and r(UUUU) TNs displayed a higher complexity with a mixture of structures, which is not yet in a full agreement with the primary NMR data. Nevertheless, all the results are significantly improved in comparison with the previous work.^12^ The presented approach is generally applicable and should further improve simulations of RNA molecules.

**Table 1.**
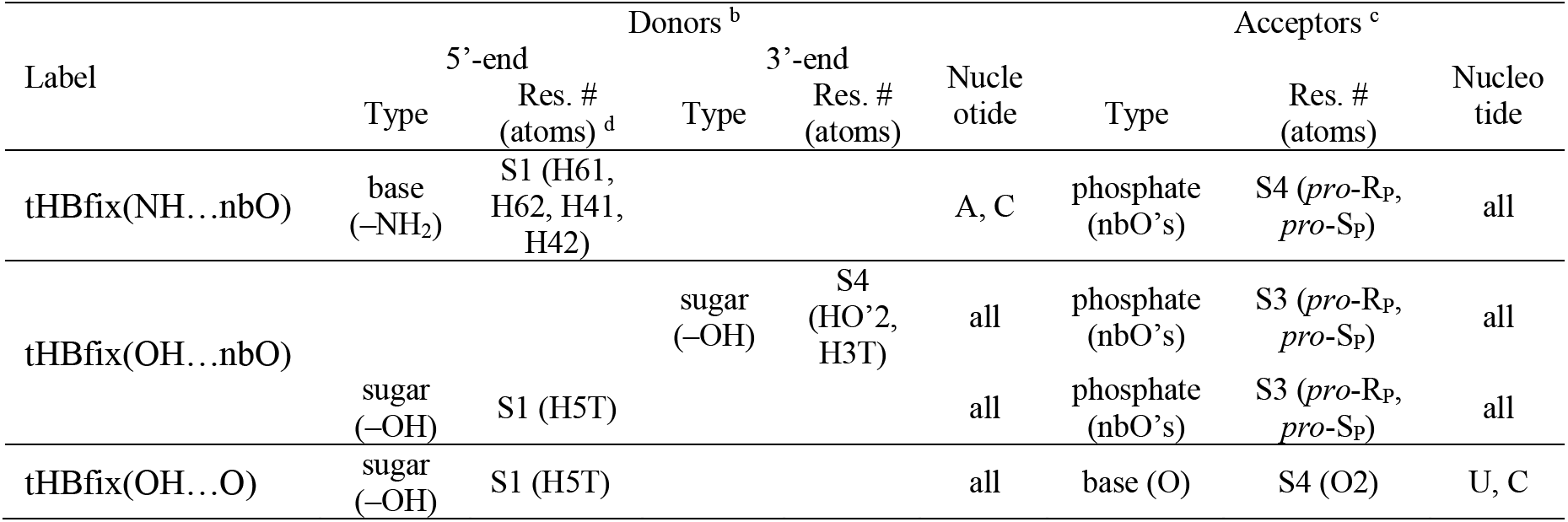
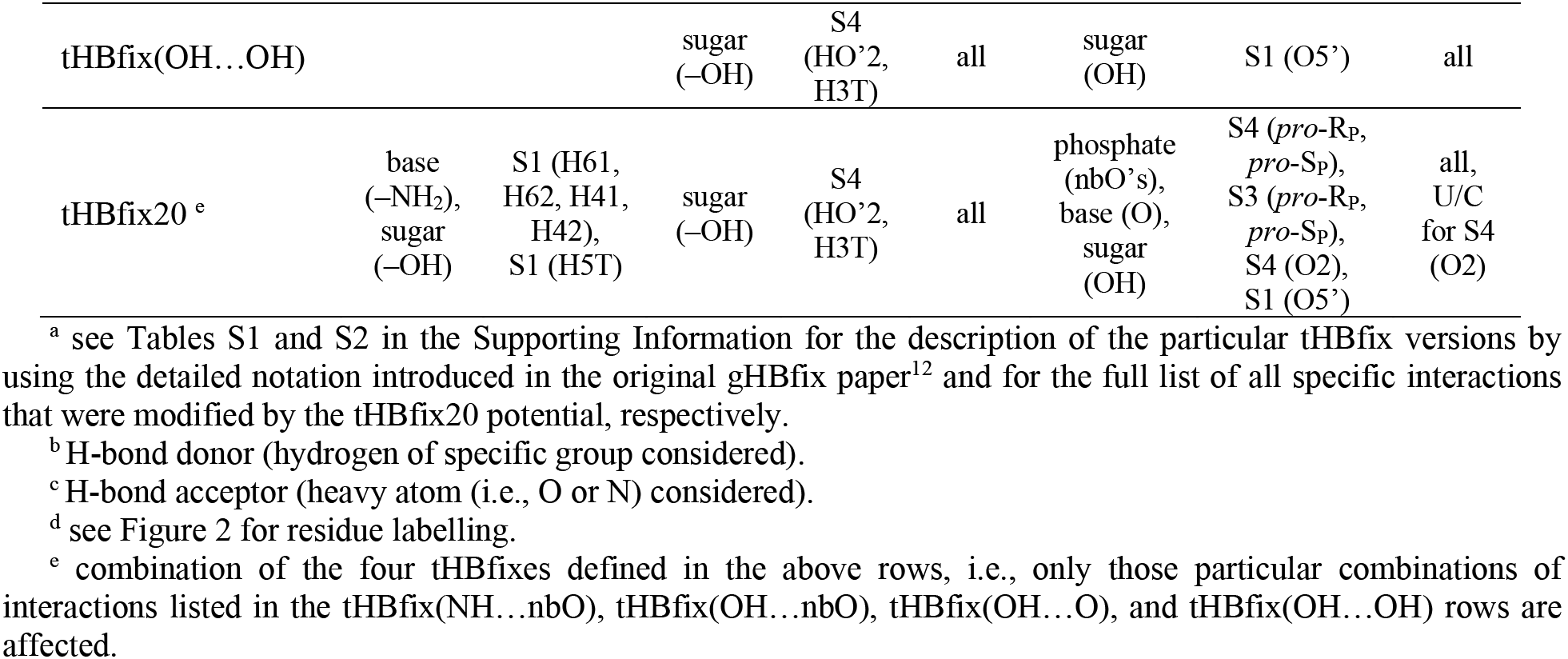
The list of terminal groups and atoms from RNA nucleotides whose interactions were modified by the different tHBfix variants.^a^

## METHODS

### Starting structures and simulation setup

The initial coordinates of r(AAAA), r(CAAU), r(CCCC), r(GACC), and r(UUUU) TNs were prepared using Nucleic Acid Builder of AmberTools14^34^ as one strand of an A-form duplex. The starting topologies and coordinates for classical MD simulations of the Sarcin-Ricin loop and the T-loop RNA motifs (see below for further details) were prepared by using the tLEaP module of AMBER 16 program package.^35^ Structures were solvated using a rectangular box with a minimum distance between box walls and solute of 12 Å, yielding ~2000 water molecules added and ~40×40×40 Å^3^ box size.

We used the standard OL3^13–16^ RNA *ff* with the vdW modification of phosphate oxygens developed by Steinbrecher et al.,^17^ where the affected dihedrals were adjusted as described elsewhere;^10, 36^ (see the Supporting Information of Ref. ^10^ for parameters) this version is abbreviated as χ_OL3CP_ henceforth. Additionally, we applied the external gHBfix potential^12^ that was shown to improve the overall *ff* performance. Most of simulations were carried out with the OPC^18^ water model and in ∼0.15 M KCl salt using the Joung−Cheatham (JC)^37^ ionic parameters. Specific tests involved TIP3P,^38^ SPC/E,^39^ and TIP4P-D^40^ water models. One test simulation of the r(GACC) TN with the TIP4P-D water was combined with charmm22 ion parameters^41^ (see the Supporting Information for other details about the simulation protocol).

### Enhanced sampling

The replica exchange solute tempering (REST2)^42^ simulations of all TNs were performed at T = 298 K with 8 replicas. Details about settings can be found in our previous work.^10^ The scaling factor (λ) values ranged from 1 to 0.601700871 and were chosen to maintain an exchange rate above 20%. The effective solute temperature ranged from 298 K to ~500 K. The hydrogen mass repartitioning^43^ with a 4-fs integration time step was used. One test simulation of r(CAAU) TN was performed at temperature 275 K corresponding to the experimental conditions.^22–24, 31^ The same λ values were applied resulting in effective solute temperature range from 275 K to ~460 K.

### HBfix-type potentials used in this work

We applied the previously introduced generalized (interaction-specific) gHBfix potential^12^ that is improving the overall performance of the χ_OL3CP_ RNA *ff*. In the majority of the simulations, we have used the basic gHBfix potential with the same parameters indicated in the earlier study^12^ as optimal (Figure 1, abbreviated here as gHBfix19, see also below). Besides that, in this work we have used several other variants of the HBfix-type refinements. We first tested additional gHBfix terms that penalized all BPh and SPh interactions in the studied systems. Later, we subtracted interactions that are established by groups from terminal nucleotides (Figure 3) and designed HBfix terms targeting only interactions that include the terminal nucleotides; these are termed as terminal-HBfix, i.e., tHBfix. The list of terminal groups and atoms from RNA nucleotides whose interactions were modified by particular tHBfix potentials are listed in Table 1 and Table S1 in the Supporting Information.

**Figure 3:**
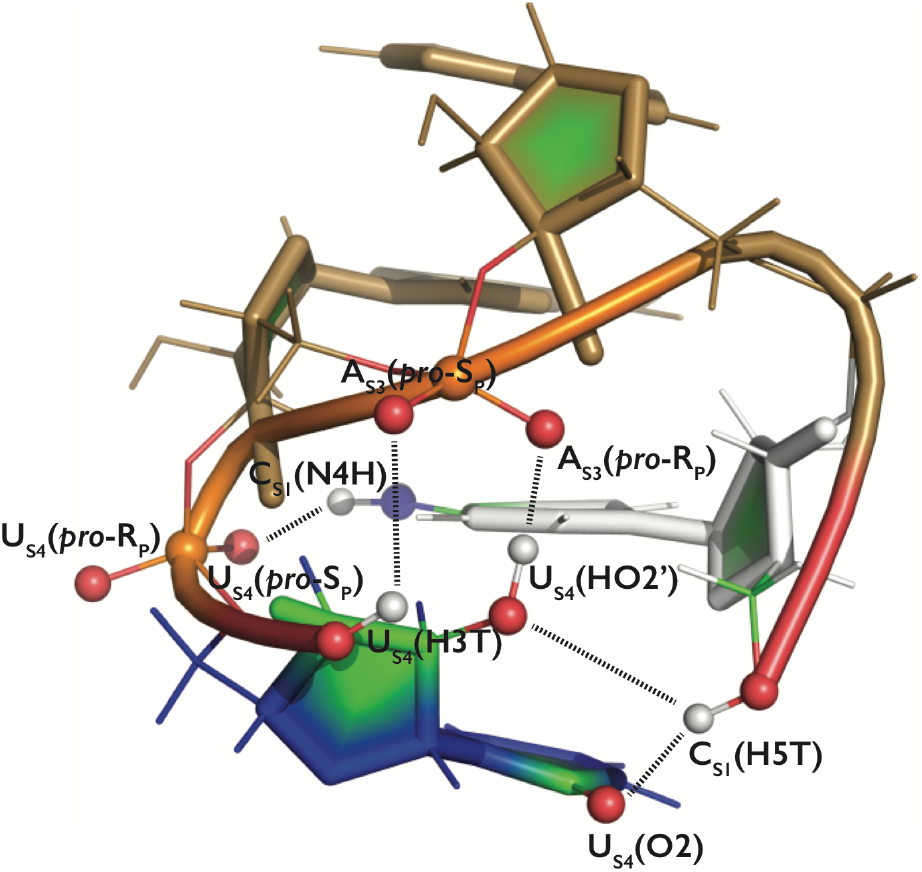
Tertiary structure of the r(CAAU) TN in the compact intercalated state. All groups responsible for the spurious H-bonds (black lines) forming BPh, SPh and sugar-base interactions are labelled and highlighted by spheres. Nucleotides are colored as in Figure 2.

### Conformational analysis

The dominant conformations sampled during REST2 simulations were identified using a cluster analysis based on an algorithm introduced by Rodriguez and Laio^44^ in combination with the *ε*RMSD^45^ (see our previous works for the detailed description of the algorithm).^10, 12^ The clustering algorithm is based on identification of the cluster center (centroid). However, this approach does not allow separation of A-RNA major and A-RNA minor conformations. Due to the definition of the cluster points and a cluster hull representing a noise spread around a given cluster,^10^ these two conformations merge into one cluster, since the transition path between them was sampled by interconversions too frequently. Thus, we performed an additional *ε*RMSD analysis, where we used representative PDB’s of the most important conformations and calculated populations of those states from MD trajectories based on simple *ε*RMSD cutoff representing a conservative definition of the conformational state (Table 2).

**Table 2:** REST2 simulations of RNA TNs using the common χ_OL3CP_ RNA *ff* combined with diverse gHBfix and tHBfix variants. All simulations used 8 replicas and were run for 10 μs (the last 7 μs were used for data analysis).^a^

| Motif | gHBfix label <sup>a</sup> | Solvent <sup>b</sup> | $\chi^2$ | Clustering [%]<br>[A-form / Loop /<br>Intercalated] | # of<br>clusters | $\epsilon\text{RMSD}$ populations [%] |
| --- | --- | --- | --- | --- | --- | --- |
|  |  |  | [<sup>3</sup>J-backbone, <sup>3</sup>J-sugar, NOE,<br>uNOE (# of violations) / **Total**] <sup>c</sup> |  |  | [A-form (major, minor) /<br>Loop (1-3, 1-4) / Intercalated] |
| r(GACC) <sup>d</sup> | gHBfix19 | OPC | 0.39, 0.50, 1.13, 0.34 (3) / **0.40** | ~74/~7/~4 | 5 | 48.0, 9.8 / 0.8, 7.0 / 2.1 |
| r(GACC) | gHBfix19 | TIP3P | 0.58, 0.48, 1.31, 0.58 (5) / **0.62** | ~57/~17/~18 | 9 | 32.5, 6.8 / 10.3, 2.7 / 11.7 |
| r(GACC) | gHBfix19 | SPC/E | 0.57, 0.66, 1.44, 0.91 (6) / **0.92** | ~60/~32/~4 | 7 | 36.6, 7.0 / 9.7, 14.4 / 1.5 |
| r(GACC) | gHBfix19_NH-N<sub>+0.75</sub> | SPC/E | 0.36, 0.41, 1.00, 0.13 (4) / **0.20** | ~84/~7/~2 | 7 | 53.3, 10.0 / 1.7, 4.0 / 2.6 |
| r(GACC) | gHBfix19 | TIP4P-D | 0.29, 0.42, 1.09, 0.00 (0) / **0.10** | ~86/~0/~3 | 5 | 67.8, 18.0 / 0.0, 0.0 / 1.1 |
| r(GACC) | gHBfix19 | TIP4P-D <sup>e</sup> | 0.32, 0.37, 1.00, 0.22 (4) / **0.28** | ~74/~8/~6 | 8 | 62.0, 7.8 / 0.0, 6.5 / 1.9 |
| r(GACC) | gHBfix19 + tHBfix20 | OPC | 0.34, 0.37, 1.11, 0.15 (3) / **0.23** | ~81/~5/~0 | 4 | 60.3, 7.2 / 0.5, 4.7 / 1.0 |
| r(GACC) | NBfix <sup>f</sup> | OPC | 1.11, 0.55, 1.08, 0.11 (5) / **0.23** | ~76/~4/~0 | 7 | 46.6, 13.2 / 1.9, 0.4 / 2.9 |
| r(CAAU) <sup>d</sup> | gHBfix19 | OPC | 0.73, 1.14, 1.81, 6.07 (28) / **5.37** | ~33/~0/~51 | 5 | 13.2, 4.6 / 0.0, 0.0 / 48.1 |
| r(CAAU) | gHBfix19 | TIP4P-D | 0.52, 1.14, 1.74, 5.04 (16) / **4.48** | ~42/~0/~51 | 4 | 19.1, 19.3 / 0.0, 0.0 / 47.8 |
| r(CAAU) | gHBfix19<sub>v2</sub>_BPh | OPC | 0.70, 0.97, 1.64, 5.18 (34) / **4.59** | ~58/~0/~22 | 8 | 18.5, 5.3 / 0.0, 0.0 / 19.9 |
| r(CAAU) | gHBfix19<sub>v2</sub>_BPh<sub>v2</sub> | OPC | 0.69, 0.95, 1.63, 4.18 (33) / **3.74** | ~55/~0/~17 | 8 | 19.5, 6.4 / 0.0, 0.0 / 13.6 |
| r(CAAU) | gHBfix19<sub>v3</sub>_BPh | OPC | 0.64, 0.81, 1.69, 2.72 (25) / **2.50** | ~64/~0/~8 | 8 | 25.8, 5.1 / 0.0, 0.0 / 6.8 |
| r(CAAU) | gHBfix19 +<br>tHBfix(NH...nbO) | SPC/E | 0.74, 1.26, 3.03, 7.67 (40) / **6.84** | ~8/~3/~69 | 5 | 3.7, 0.9 / 0.0, 0.0 / 40.3 |
| r(CAAU) | gHBfix19 +<br>tHBfix(NH...nbO) <sup>g</sup> | SPC/E | 1.17, 1.09, 5.70, 8.45 (36) / **7.75** | ~0/~0/~88 | 3 | 0.8, 0.7 / 0.0, 0.0 / 62.5 |
| r(CAAU) | gHBfix19 +<br>tHBfix(NH...nbO) | OPC | 0.56, 1.01, 1.33, 4.09 (33) / **3.63** | ~63/~0/~19 | 6 | 19.9, 8.5 / 0.0, 0.0 / 16.0 |
| r(CAAU) | gHBfix19 +<br>tHBfix(NH...nbO) <sup>g</sup> | OPC | 0.54, 0.96, 1.25, 3.82 (34) / **3.40** | ~61/~0/~17 | 9 | 22.3, 8.4 / 0.0, 0.0 / 13.9 |
| r(CAAU) | gHBfix19 + tHBfix20 | OPC | 0.49, 0.80, 1.40, 2.22 (29) / **2.05** | ~74/~0/~7 | 8 | 29.1, 6.8 / 0.0, 0.0 / 5.1 |
| r(CAAU) | gHBfix19 + tHBfix20 <sup>h</sup> | OPC | 0.49, 0.88, 1.37, 2.56 (32) / **2.34** | ~68/~0/~3 | 8 | 31.1, 5.5 / 0.0, 0.0 / 2.3 |
| r(AAAA) <sup>d</sup> | gHBfix19 | OPC | 0.66, 1.23, 1.08, 1.14 (12) / **1.11** | ~30/~0/~10 | 17 | 12.1, 4.4 / 0.0, 0.0 / 8.1 |
| r(AAAA) | gHBfix19 | TIP4P-D | 0.69, 1.12, 1.03, 2.22 (11) / **1.96** | ~29/~0/~32 | 13 | 14.6, 6.8 / 0.0, 0.0 / 26.8 |
| r(AAAA) | gHBfix19 +<br>tHBfix(NH...nbO) | OPC | 0.67, 1.38, 1.20, 1.02 (11) / **1.04** | ~38/~0/~5 | 16 | 12.1, 4.5 / 0.0, 0.0 / 1.7 |
| r(AAAA) | gHBfix19 + tHBfix20 | OPC | 0.68, 1.48, 1.28, 1.06 (11) / **1.08** | ~32/~0/~1 | 18 | 12.5, 4.5 / 0.0, 0.0 / 1.2 |
| r(CCCC)<sup>d</sup> | gHBfix19 | OPC | 0.44, 0.68, 1.54, 3.22 (11) / **2.83** | ~55/~0/~40 | 2 | 39.1, 5.5 / 0.0, 0.0 / 37.1 |
| r(CCCC) | gHBfix19 | TIP4P-D | 0.45, 0.53, 1.73, 3.72 (10) / **3.26** | ~49/~0/~48 | 3 | 41.0, 6.4 / 0.0, 0.0 / 46.6 |
| r(CCCC) | gHBfix19 +<br>tHBfix(NH...nbO) | OPC | 0.31, 0.46, 1.14, 1.30 (9) / **1.20** | ~74/~0/~20 | 5 | 52.2, 7.8 / 0.0, 0.0 / 26.4 |
| r(CCCC) | gHBfix19 + tHBfix20 | OPC | 0.25, 0.38, 1.22, 0.50 (6) / **0.55** | ~85/~0/~8 | 5 | 63.3, 7.1 / 0.0, 0.0 / 6.1 |
| r(UUUU)<sup>d</sup> | gHBfix19 | OPC | 0.53, 0.87, 3.53, 1.65 (19) / **1.63** | ~5/~0/~3 | 10 | 6.6, 1.7 / 0.0, 0.0 / 3.8 |
| r(UUUU) | gHBfix19 | TIP4P-D | 0.52, 0.72, 3.85, 1.52 (16) / **1.51** | ~25/~0/~4 | 13 | 14.0, 6.1 / 0.0, 0.0 / 3.8 |
| r(UUUU) | gHBfix19 + tHBfix20 | OPC | 0.52, 0.97, 3.46, 1.86 (17) / **1.81** | ~14/~0/~0 | 10 | 6.4, 1.5 / 0.0, 0.0 / 1.9 |

<sup>a</sup> see Table S1 in the SI for the detailed definition of the tested variants of the gHBfix and tHBfix potentials.
<sup>b</sup> 0.15 M KCl, JC ion parameters.<sup>37</sup>
<sup>c</sup> $\chi^2$ values were obtained by comparing calculated and experimental backbone <sup>3</sup>J scalar couplings, sugar <sup>3</sup>J scalar couplings, nuclear Overhauser effect intensities (NOEs), and the absence of specific peaks in NOE spectroscopy (uNOEs, number of violations is also reported). Note that we identified some potential uncertainty in the experimental datasets, which resulted in higher $\chi^2$ values and lowered the agreement between simulations and the experiment for r(CCCC) and r(UUUU) TNs (see Methods for details).
<sup>d</sup> data taken from the preceding gHBfix paper.<sup>12</sup>
<sup>e</sup> simulation run with charmm22 ion parameters.<sup>41</sup>
<sup>f</sup> NBfix *ff* reparameterization was prepared in a way to be comparable with the combined gHBfix19 + tHBfix20 potential (see Table S4 in the Supporting Information for details).
<sup>g</sup> extended variant of tHBfix(NH...nbO) between –NH<sub>2</sub> group from Cs<sub>1</sub> (5'-end) and nbO atoms of *all phosphates*, i.e., not only nbOs from the terminal U<sub>s4</sub> nucleotide.
<sup>h</sup> REST2 simulation run with unbiased replica shifted from 298 K to 275 K (to mimic the experimental temperature).

The convergence was checked by bootstrapping^46^ using the recently introduced implementation.^10, 12^ Errors were estimated by resampling of the time blocks of the whole set of replicas and subsequent resampling of the coordinate-following replicas for the purpose of their demultiplexing to follow Hamiltonians in order to obtain a final resampled population at the reference unbiased Hamiltonian state (see Figure S1 in the Supporting Information of Ref. ^12^ for more details). Figure S1 in the Supporting Information shows that the major conformers are uniformly represented within all replicas from the particular REST2 simulations.

### Stacking analysis

We implemented an in-house code in order to probe the effect of stacking on the *syn/anti* balance of nucleobases. All heavy atoms of each nucleobase are fitted by a plane,^47^ projected into this plane and transformed to a convex planar polygon.^48^ For each pair of polygons, the program quantifies their geometrical overlap as follows: A part of the second polygon in the stacking distance (from 3 to 4) is projected into the plane of the first polygon and the area of calculated intersection between both polygons is measured. The calculated values of the geometrical overlap for consecutive nucleobases lie between zero and one (in relative values), where zero means no overlap and one is the maximum overlap.

### Comparison with the experiment

The conformational ensembles obtained from MD simulations were compared with solution experiments.^22–24^ We used and analyzed separately four NMR observables, i.e., (i) backbone ^3^J scalar couplings, (ii) sugar ^3^J scalar couplings, (iii) nuclear Overhauser effect intensities (NOEs), and (iv) the absence of specific peaks in NOE spectroscopy (uNOEs). Their combination provided the total χ^2^ value for each REST2 simulation. The lower χ^2^ value, the better agreement between the experimental data and the conformational ensemble from the particular simulation was achieved. Note that all χ^2^ values below 1 indicate good agreement with the experiment and, considering the experimental error, it is not straightforward to decide among two simulations with χ^2^ < 1, which of them agrees better with the NMR datasets. However, the above-mentioned rule of thumb of χ^2^ < 1 can be applied primarily to the χ^2^ contributions coming from ^3^J scalar couplings and NOEs as they rigorously follow statistical χ^2^ distribution. In turn, the component coming from uNOEs rather qualitatively indicates violations of the NOEs data for particular contacts, so the interpretation of this χ^2^ value is not straightforward. Thus, although simulations with *total* χ^2^ < 1 might be considered to be in agreement with the experimental data, one should always check also the *individual* χ^2^ components corresponding to the particular NMR observables.

We further identified that the signal range for some NOEs, i.e., C2(H6)…C2(H5’’), U2(H3’)…U2(H5’), and U3(H3’)…U3(H5’), was rarely populated during MD simulations. The experimental NOEs would thus indicate some rather atypical RNA backbone conformations which might also indicate some uncertainty in the experimental data. Those signals provided higher χ^2^ values (for all simulations of the particular TN sequence) and thus, lowered the agreement between simulations and the experiment for r(CCCC) and r(UUUU) TNs (Table 2).

NOEs and uNOEs were obtained from MD simulations as averages over the N samples, i.e., 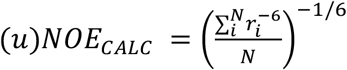. The calculated NOEs were directly compared against experimental values. Each uNOE signal was considered as violated when the calculated value from the simulation was shorter than the assigned one from the experiment. ^3^J scalar couplings were obtained from the Karplus relationships (see Supporting Information of Ref. ^30^ for the details and experimental datasets).

## RESULTS AND DISCUSSION

In our preceding work we demonstrated that performance of the basic OL3 AMBER RNA *ff*^13, 15–17, 42^ is improved by using the gHBfix 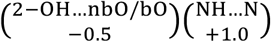 variant,12 abbreviated in the present paper as gHBfix19, see Table S1. This gHBfix version weakens all SPh H-bonds by 0.5 kcal/mol and supports all base-base –NH…N– H-bonds by 1.0 kcal/mol (Figure 1). These parameters were derived to be combined with the OPC water model and the OL3 *ff* has been used with modified phosphate vdW parameters (χ_OL3CP_, see Methods). In the present work we used the χ_OL3CP_ basic RNA *ff* as the starting version and tried to further improve its performance. We focused on five RNA TNs, i.e., r(AAAA), r(CAAU), r(CCCC), r(GACC), and r(UUUU), where highly accurate experimental data are available.^22–24^ The application of the basic gHBfix19 potential leads to visible but only partial improvement in structural description of three TNs.^12^ More specifically, population of intercalated structures still remained significant for r(CAAU), r(AAAA), and r(CCCC) TNs.^12^ Thus we tried additional means to improve the TN simulations.

### Weakening of BPh interactions in the framework of gHBfix improves TN simulations

In our previous work,^12^ we found out that sequences with higher propensity to form the intercalated structures, i.e., r(CAAU), r(AAAA), and r(CCCC), possess C or A as the 5’-terminal nucleotide. These nucleotides can form type-7 BPh (7BPh) interaction^49^ in the intercalated state (Figure 4). BPh interactions were previously reported to be generally overpopulated in AMBER simulations of unfolded states, contributing to spurious compaction of RNA single strands and thus hindering folding of RNA tetraloops.^10^ We have suggested that their weakening could suppress the spurious intercalated structures of TNs.^12^ However, native BPh interactions are common in many folded RNA structures^49^ and thus tuning of these interactions was not attempted in Ref. ^12^.

**Figure 4:**
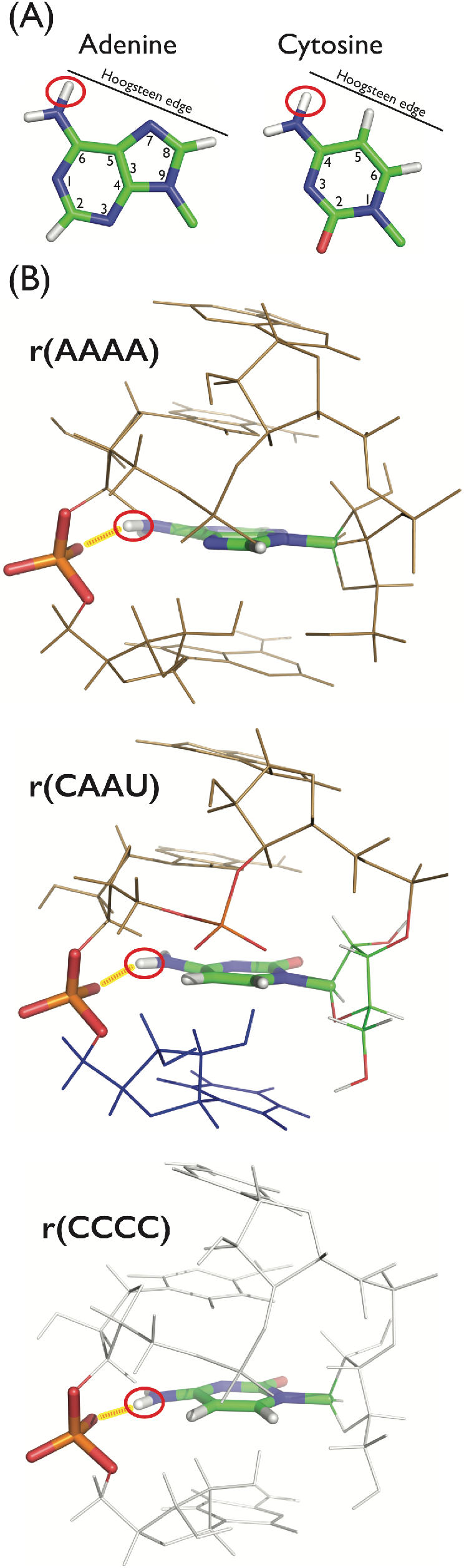
The spurious intercalated state is highly populated during MD simulations of TNs containing C or A as the 5’-terminal nucleotide. (A) The 7BPh interaction^49^ is formed by nucleotides that possess the amino group at the Hoogsteen edge, i.e., N6H and N4H for A and C, respectively (red circle). (B) The detail of the 7BPh interaction^49^ in r(AAAA), r(CAAU), and r(CCCC) TNs. The spurious H-bond between the amino group and nbOs of the nucleotide at the 3’-end is highlighted (red line in yellow background).

In the present work we used the r(CAAU) TN as the initial testing system, as this TN has the intercalated structure more populated than the native A-form even with the gHBfix19; see Table 6 in Ref. ^12^. We have prepared three new gHBfixes, i.e., gHBfix19_v2__BPh, gHBfix19_v2__BPh_v2_, and gHBfix19_v3__BPh, where the gHBfix19 is combined with either 0.5 (BPh) or 1.0 (BPh_v2_) kcal/mol penalty of all possible BPh interactions (see Table S1 for full details; the description of SPh interactions in gHBfix19_v2_ is marginally different from the gHBfix19 which should, however, have no practical effect on the simulations). In addition, in the gHBfix19_v3__BPh version, we further weakened (from 0.5 to 1.0 kcal/mol) the SPh interactions. We observed that weakening of BPh interactions is progressively improving the simulation outcome as the higher penalty to –NH…nbO– contacts decreased the population of intercalated structures more efficiently and, consequently, provided lower χ^2^ values (Table 2). Interestingly, weakening of BPh interactions allowed us to increase penalty to the SPh interactions. With the gHBfix19 setting the SPh interactions still appear to be overstabilized but their additional weakening (without penalizing BPh interactions) would eliminate occurrence of the A-RNA minor conformation from the structural ensemble.^12^ A-RNA minor conformation is an auxiliary A-RNA conformation of the TN accompanying the major A-RNA form and, according to the experiments, it should be populated to a certain extent.^22^ The best result among those three simulations was obtained with the gHBfix19_v3__BPh version, where all possible BPh interactions were penalized by 0.5 kcal/mol (χ^2^ value of 2.50, Table 2). The small penalty sufficient for the BPh interactions is promising since their large weakening could destabilize structures of folded RNAs, where BPh contacts are common and highly conserved.^49^

In order to explicitly test the effect of weakening of BPh interactions for structural description of important RNA motifs, we used two systems with multiple conserved BPh interactions, namely the Sarcin-Ricin loop RNA motif (SRL, PDB ID 3DW4^50^), a highly conserved RNA motif which was originally found in helix 95 of domain VI of the large ribosomal subunit,^51^ and the T-loop RNA motif, a frequently occurring five-nucleotide hairpin motif assuming U-turn-like structures.^52^ The ucuUGGAAcaga 12-mer sequence was used as the T-loop model system for testing (extracted from the PDB ID 1JJ2,^53^ residues 310 – 321). T-loops and especially the SRL motif are established testing systems for RNA *ff* validations because they contain an intricate network of non-canonical base pairs and BPh interactions, which is complemented by very complex backbone conformation. We performed standard MD simulations (few μs-long timescales) using the χ_OL3CP_ RNA *ff* with the gHBfix19_v3__BPh version and observed that the partial weakening of BPh contacts is not *a priori* detrimental, because the overall fold and all key interactions within those motifs were maintained (Figures S2 and S3 in the Supporting Information). However, we cannot rule out possible side effects on longer time scales and/or for other systems. Much broader testing considering weakening of all BPh interactions would be required before introducing a *ff* version with all BPh contacts been weakened using the gHBfix *ff* term.

### Weakening of specific interactions formed by terminal groups is comparable with tuning of all BPh contacts

Although BPh interactions appear to be over-stabilized within the current basic RNA AMBER *ff*, their general penalization could introduce side-effects. Despite that the above simulations on SRL and T-loop motifs were not affected, a complete testing of weakened BPh interactions would be a major challenge. Thus, we considered construction of terminal-nucleotide-specific HBfixes (tHBfixes), i.e. HBfix-type modifications where one or both interacting atoms belong to the terminal residues. This should allow to tune simulations of TNs while not introducing side effects for the other systems. The RNA sequences most prone to form intercalated structures, i.e., r(CAAU), r(AAAA), and r(CCCC), contain –NH_2_ group at the Hoogsteen edge of the nucleotide (either C_S1_ or A_S1_) at the 5’-end. This amino group is forming 7BPh interaction^49^ (Figure 4) with nbOs of the other terminal nucleotide at the 3’-end. Thus, a penalty to this specific interaction between groups from nucleotides on 5’- and 3’-ends should not introduce any side effects for other RNA structures as it is changing interactions only between two terminal nucleotides. We designed the gHBfix19 + tHBfix(NH…nbO) variant (Table 1), which is composed of the basic gHBfix19^12^ extended by tHBfix penalizing interactions between –NH_2_ group from either C_S1_/A_S1_ (5‘-end) and both nbOs of the 3‘-end nucleotide by 1.0 kcal/mol (Table S1). We firstly tested the effect on r(CAAU) TN and observed that the agreement with experimental data is improved (χ^2^ value of 3.63 is slightly worse than χ^2^ of 2.50 from the simulation with gHBfix19_v3__BPh applied to all BPh contacts, while the basic gHBfix19 value is 5.37, see Table 2). However, the population of intercalated structures is still significant (~16%, Table 2). Thus, the introduced gHBfix19 + tHBfix(NH…nbO) potential leads to clear improvement, but the extra 1.0 kcal/mol penalty to the single H-bond contact is still not sufficient to fully eliminate the occurrence of the intercalated state of r(CAAU) TN.

We also probed the effect of the extended gHBfix19 + tHBfix(NH…nbO) potential to all phosphates. Thus we penalized (by 1.0 kcal/mol) interactions between the –NH_2_ group from the terminal C_S1_ (5‘-end) and nbOs from all nucleotides of the r(CAAU) TN. We observed that the resulting conformational ensembles of these two different implementations of the gHBfix19 + tHBfix(NH…nbO) potential (i.e., biasing BPh interactions formed by nbOs of the phosphate from terminal nucleotide only and from all phosphates) are comparable within the limit of sampling (Table 2). Such a result was also confirmed when using the SPC/E water model instead of OPC (Table 2). We, however, tentatively propose that the second extended variant of the tHBfix(NH…nbO) potential (involving nbOs from all phosphates) might be useful for tuning simulations of larger RNA molecules, where different spurious (intercalated) states stabilized by BPh interactions involving the –NH_2_ group from the 5’-end terminal nucleotide and nbOs from phosphates of different internal nucleotides might occur.

### Intercalated states can be suppressed by penalizing SPh interactions formed by terminal –OH groups

Closer inspection of the intercalated state (Figures 2 and 3) revealed intricate network of SPh contacts, where the majority of them involve terminal –OH groups. Although these are already destabilized within the basic gHBfix19 by 0.5 kcal/mol, it appears that such destabilization is insufficient to fully correct the behavior of terminal –OH groups during MD simulations. Since the terminal –OH groups contain specific parameters within current *ff*s, e.g., they possess different electrostatic and vdW parameters compared to general –OH groups, it is entirely reasonable to treat their interaction separately from interactions established by the other–OH groups. Further, adding stronger destabilization for these specific contacts rather than penalizing all SPh interactions should be significantly less prone towards introducing undesirable side effects, e.g., elimination of A-RNA minor conformers, as discussed previously.^12^ Thus, we designed the gHBfix19 + tHBfix20 variant for the CAAU sequence, where the additional tHBfix terms are destabilizing the following H-bonds by 1.0 kcal/mol: (i) the C_S1_(N4H)…U_S4_(*pro*-R_P_/*pro*-S_P_) BPh, (ii) the U_S4_(2’-OH/3’-OH)…A_S3_(*pro*-R_P_/*pro*-S_P_) SPh, (iii) the C_S1_(5’-OH)…A_S3_(*pro*-R_P_/*pro*-S_P_) SPh, (iv) the C_S1_(5’-OH)…U_S4_(O2) sugar-base H-bond, and (v) the U_S4_(2’-OH/3’-OH)…C_S1_(O5’) sugar-sugar interactions (Table 1 and Table S1 in the Supporting Information). The REST2 simulation revealed the best agreement with the experiment for r(CAAU) so far (the obtained χ^2^ value of 2.05 is lowest so far, see Table 2). Thus, the structural dynamics of the r(CAAU) during REST2 simulations is significantly improved and the population of intercalated structures is reasonably low (~5%, Table 2).

### The gHBfix19 + tHBfix20 potential improves the structural dynamics of other TNs

We have carried out REST2 simulations with the gHBfix19 + tHBfix20 for the remaining four TNs. The newly designed potential significantly improves the structural dynamics of the r(CCCC) TN (χ^2^ value of 0.55, Table 2), since the agreement between all calculated and experimental observables, i.e., ^3^J-backbone couplings, ^3^J-sugar couplings, NOE and uNOE signals, is better (Table 2). The huge decrease of the χ^2^ value coming from uNOE signals correlates with the decreased population of the spurious intercalated structure (from ~37% in simulation with the basic gHBfix19^12^ to ~6%). For the r(GACC) TN, the calculated χ^2^ value remains low (0.23) and is comparable with the value from the simulation with the basic gHBfix19^12^ (χ^2^ value of 0.40, Table 2). It is not possible to decide, which simulations agrees better with the experiment since all simulations with χ^2^ value below 1 are satisfactory (see Methods).

The results for the remaining two TNs, i.e., r(AAAA) and r(UUUU), are ambiguous. Importantly, gHBfix19 + tHBfix20 eliminates population of intercalated structures within conformational ensembles of both TNs (to 1.2% and 1.9% for r(AAAA) and r(UUUU), respectively). However, total χ^2^ values remain still rather high (1.08 and 1.81 for r(AAAA) and r(UUUU), respectively) indicating that the populations of states during MD simulations are not fully consistent with the experimental ensemble (Table 2). However, it is difficult to exactly pinpoint the source of the difference. The χ^2^ value of 1.81 for r(UUUU) TN is even marginally higher than the one from the simulation with the basic gHBfix19 (1.63).^12^ Conformational ensembles of r(AAAA) and r(UUUU) REST2 simulations show that the major clusters are different from the canonical A-RNA major/minor conformers. The r(UUUU) TN is expected to be highly dynamical and to sample diverse conformations. We identified that 1-3/2-4 stacked structures are preferred during the REST2 simulation (clusters 1, 4 and 5 with total ~25% populations, with ~35% conformations unassigned; see Figure S4 in the Supporting Information). These states (typically with *syn* conformation of at least one U) appear to cause errors in NOEs and uNOEs data.

r(AAAA) TN preferred the four-stack state during simulations, but we noticed that two out of four nucleotides often favored *syn* conformation of χ dihedral (Figure S5 in the Supporting Information). The A_S1_ nucleotide at the 5’-end revealed higher propensity to sample *syn* conformation. Although it can be explained by the *syn-*specific A_S1_(5’-OH)…A_S1_(N3) H-bond,^54^ it is possible that the *syn* state is overpopulated. Interestingly, in Ref. ^27^ the combination of MD simulations with NMR data for nucleotides and dinucleotides suggested the need for increasing stabilization of the *anti* conformation for A nucleotides in the *ff*. Similar indication was obtained from another fitting based on the same experimental data as used here^55^ (see https://github.com/bussilab/ff-fitting-tools/blob/master/Analysis.ipynb). In other words, the large tendency of the A nucleotide to sample *syn* conformation could indicate that its *syn/anti* balance is not fully correct in the χ_OL3CP_ RNA *ff*, suggesting a possible modification of the dihedral potential. However, the likely spurious preference for *syn* orientation in r(AAAA) simulations might be also caused by not enough accurate description of the stacking interactions. Stacking is suspected to be overestimated within current RNA *ff*s,^56–58^ albeit the OPC water model might partly reduce such overestimation.^19, 59^

We performed stacking overlap analysis with an in-house program and probed the possible effect of stacking on the *syn/anti* balance of r(AAAA) TN by calculating and quantifying the geometrical overlap of the stacked nucleobases in their various *syn/anti* orientations, as sampled in the simulations (see Methods for details). Initially, we extracted A-form-like states from the conformational ensemble, i.e., states, where consecutive nucleotides are stacked on top of each other in the correct order (A_S1_-A_S2_-A_S3_-A_S4_). Among 16 possible combinations of *anti/syn* states for 4 nucleotides, all *anti* structures were indeed the most populated (~49%), followed by *syn-anti-anti-anti* (~14%), *anti-anti-anti-syn* (~12%), *syn-anti-syn-anti* (~9%) and *anti-anti-syn-anti* (~7%) conformational states. Remaining combinations of glycosidic orientations were marginally populated (below 3%). Subsequently, we analyzed the geometrical overlap of the consecutively stacked nucleobases, i.e., we separately analyzed A_S1_-A_S2_ (5’-end), A_S2_-A_S3_ (middle), and A_S3_-A_S4_ (3’-end) stacking pairs, in their various *syn/anti* orientations. The largest stacking overlap is achieved with *anti/anti* combination at the 5’-end, but *syn/syn* and *syn/anti* orientations have the largest stacking overlap in the middle step and at the 3’end, respectively (Table S3 in the Supporting Information). Based on this simple analysis, we suggest a possibility that overestimated stacking interaction is pushing the A_S3_ nucleotide to sample *syn* state more frequently during MD simulations. Nevertheless, the potentially excessive population of *syn* states in simulations of r(AAAA) may be a complex problem caused by a combination of different factors which requires further investigations.

### gHBfix and tHBfix combined with different solvent models

The OPC water model^18^ is nowadays widely used for enhanced-sampling simulations of simple RNA motifs^10, 12, 26–28, 30, 60^ because it somewhat reduces the excessive formation of SPh contacts,^19^ especially in combination with the revised phosphate parameters.^17^ On the other hand, the OPC water is suboptimal for G-quadruplex simulations.^61, 62^ It indicates that it may not represent a universal solution for MD simulations of all nucleic acids, since desirable weakening of spurious interactions can sometimes be accompanied by undesirable destabilization of native interactions. Thus, we investigated effects of the gHBfix correction to the RNA *ff* in combination with different solvent models.

We firstly tested the TIP3P water model^38, 63^ during REST2 simulations of r(GACC) with the gHBfix19 potential.^12^ The population of intercalated structures is significantly increased (~12%, Table 2) and the χ^2^ value indicating the disagreement with the experimental data rises up to 0.62. This result is similar to simulations with the OPC water without any gHBfix (χ^2^ value of 0.75, Table 3 in Ref. ^12^). In addition, we observed enhanced probability to form loop-like structures, where G_S1_ established the Watson-Crick base pairs with C_S3_ and C_S4_ forming 1-3 loops (~10% population) and 1-4 loops (~3% population), respectively (Table 2, Figure S6 in the Supporting Information).

Next, we performed gHBfix19 simulation of the r(GACC) TN with the SPC/E water model,^39, 64^ which is often used in simulations of protein-RNA complexes.^11^ The simulation revealed higher total χ^2^ value of 0.92, which was mostly caused by the difference between predicted and observed NOE distances and by the fact that the MD simulation predicts signals that are not observed by the experiment (unobserved NOE signals, Table 2). The increased χ^2^ value of uNOEs (from 0.34 to 0.91 in OPC and SPC/E simulation, respectively) could be related to increase of populations of loop-like states from ~7% to ~32% in OPC and SPC/E simulation, respectively. Interestingly, the population of r(GACC) intercalated structures is low (~2%), comparable to the OPC data (Table 2). Thus, we designed a new gHBfix19_NH-N_+0.75_, where we slightly weakened base-base interactions by decreasing the bias (from 1.0 kcal/mol to 0.75 kcal/mol) to all possible –NH…N– contacts (Table S1). Population of loop-like states dropped to ~7% (from ~32% in the simulation with the basic gHBfix19 potential, Table 2). Then we tested the effect of SPC/E water model in two simulations of r(CAAU) TN using the combined gHBfix19 + tHBfix(NH…nbO) terms. Unfortunately, population of intercalated states significantly increased (reaching more than 60%), which correlates with high total χ^2^ values of 6.84 and 7.75 (Table 2). Thus, SPC/E water model revealed comparable results with the OPC model for r(GACC) TN upon minor modification of gHBfix19 but results for r(CAAU) suggest that the SPC/E performance could be system-dependent. Finding of suitable gHBfix + tHBfix parameters with the SPC/E water model would require further work.

Finally, we also tested the TIP4P-D water model, which was initially derived for simulations of disordered proteins^40^ and was also recently suggested for RNA simulations.^32^ Initially, we tested the TIP4P-D model during REST2 simulation of the r(GACC) TN using two different ion parameters, i.e., JC^37^ and charmm22.^41^ Results from both REST2 simulations of the r(GACC) TN revealed good agreement with the experiment (Table 2) and are comparable to the simulation with the OPC water model. Subsequently, we tested the abovementioned combination of the basic gHBfix19, TIP4P-D water and JC ions on the other TN’s from the set. We observed that TIP4P-D water model is providing comparable results with the OPC model considering the r(CAAU), r(CCCC), and r(UUUU) TNs (the χ^2^ value is slightly lower for r(CAAU) TN and slightly higher for both r(CCCC) and r(UUUU) TNs, Table 2). However, r(AAAA) REST2 simulations showed that the agreement with the experiment is much worse in comparison with the OPC water model and the amount of spurious intercalated states is significantly increased (from ~8% to ~27%, Table 2). Thus, it appears that the performance of the TIP4P-D water model is typically on-par with the OPC water, but for some systems its behavior may deteriorate (as observed also for the SPC/E model). The reason causing the difference between SPC/E and TIP4P-D water models for the r(AAAA) is not known and this point will require further research. Some role of the ions cannot be excluded^33^ but full testing is beyond the scope of this work due to the high computer demands.

In conclusion, although our results are far from providing an extensive testing of the water models it seems that usage of three-point water models, e.g., TIP3P and SPC/E, instead of OPC in TN simulations increases the population of loop-like and/or intercalated states. However, it cannot be ruled out that it would be possible to find gHBfix and tHBfix settings which provide better results also with these water models. The gHBfix settings, however, would have to be different from those used with OPC. The simulation outcome with the TIP4P-D water was generally comparable with the OPC water model, except of the rAAAA TN. We note that for this water model we did not test the tHBfix parameters though we assume that the effect of the tHBfix could be similar to the OPC water model. The data support the view that the OPC model may be optimal for TN simulations, though transferability of its performance to other nucleic acids systems is not a priori guaranteed.^61^ The results confirm sensitivity of nucleic acids simulations to water models.^4, 19, 61, 65, 66^ This complicates tuning of nucleic acids *ff*s, especially when refining the non-bonded terms. Massive testing would be required to fully clarify the interrelation between the water models and the HBfix-type corrections, which is however, out of scope of this work.

### Reference temperature does not affect the agreement between computed and experimental results

The NMR datasets of TN motifs were measured at 275 K,^22–24^ whereas we calculate the NMR observables from MD simulations using 298 K reference replica. Thus, we explicitly checked if and how the difference in temperature is affecting the comparison. We performed additional simulation using the newly designed gHBfix19 + tHBfix20 potential with the unbiased (reference) replica shifted to 275 K (see Methods section for the details). The outcome is comparable within limits of sampling to the 298 K simulation (Table 2). Thus, we expect that TN conformational ensembles are equivalent at both temperatures and the results from experiments and computations can be compared despite the temperature difference.

### Note about the usage of reweighting procedures for the newly-designed gHBfix potentials

When working on the gHBfix19 and tHBfix20 parameters, we usually tested the parameters on a grid containing energy values of –1.0, –0.5, +0.0, +0.5 or +1.0 kcal/mol and only optimized a limited number of parameters at the same time.^12^ However, the simulations do indicate that in some cases finer optimization could have visible influence on the results. We made an initial attempt to refine simultaneously all the parameters of the gHBfix potential by using the reweighting scheme proposed in Ref ^55^. We used the set of fourteen available REST2 simulations of r(GACC) TN from our previous work, where different bias to base-base and SPh interactions was applied.^12^ However, the outcome from reweighting showed us that the protocol using just one system (albeit with broad set of simulations with different parameters) with too many parameters is prone to suggest the optimized values of gHBfix parameters too far from the initial *ff*, which might be explained by overfitting to one particular system. For example, the reweighting scheme using only simulations of the r(GACC) TN motif suggested to significantly further penalize most BPh and sugar-base interactions. Thus, in order to use the reweighting scheme to simultaneously tune a larger number of parameters, one needs to also add data from converged simulations of other motifs, where these interactions are present in different contexts. This will be addressed in future studies.

### The possibility to reproduce the effect of the combined gHBfix19 and tHBfix20 potential by the NBfix approach

In principle, similar effect as obtained by the HBfix-type of potentials could be achieved by modification of the pairwise vdW parameters via breakage of the combination (mixing) rules, i.e., the so-called nonbonded fix (NBfix) approach; see recent review^67^ for details. However, as we discussed earlier,^12^ use of the HBfix-type approach is more straightforward and brings some advantages. To tune all possible H-bonds by the NBfix approach is a demanding work with huge dimensionality, as it is necessary to create new atom types to differentiate atoms on different residues with same atom types and different partial charges. In addition, as discussed by us in Ref. ^12^, the NBfix approach affects both interaction energy and optimal geometry, so its applicability may be limited by geometrical constraints. Thus, the HBfix terms (e.g., gHBfix, tHBfix) are more flexible for fine-tuning of the non-bonded interactions with respect to the NBfix approach in pair-specific treatment of H-bonds and interatomic contacts. In the preceding study we have nevertheless published a set of NBfix parameters that have similar effect as the gHBfix19 potential.^12^ In practice, the NBfix parameters could be derived a posteriori after finding the suitable gHBfix version, since the gHBfix allows more direct tuning of the H-bonds. Nevertheless, it is probable that both approaches will be combined in future for tuning of *ff*s.

In the present work we also derived the NBfix parameters to reproduce the interaction energy curves obtained with the gHBfix19 and tHBfix20 terms in the same way as it is discussed in Ref. ^12^. Details are provided in the Supporting Information together with the NBfix parameters which can be used in simulations as an alternative to the gHBfix19 + tHBfix20 parameter set. Note that to emulate the effect of the tHBfix20 potential by the NBfix terms, one needs to introduce new atom types that allow to differentiate between interactions formed by groups from terminal and other residues (sharing the same atom types) in order to apply different parameters. We subsequently tested the NBfix correction, which modified base-base, SPh and BPh interactions (Table S4 in the Supporting Information), in a REST2 simulation of GACC TN. We observed that the NBfix simulation and the original simulation modified by the combined gHBfix19 + tHBfix20 potential are comparable. Both revealed similar χ^2^ value of 0.23 and populations of major conformers within both ensembles were only marginally different (Table 2). Due to the computer demands we did not make testing for the remaining TNs.

## CONCLUDING REMARKS

Performance of pair-additive biomolecular *ffs* is determined by a wide range of parameters. Although the most common approach to refine biomolecular *ffs* is tuning of the dihedral potentials, it is becoming clear that refinement of RNA *ff*s requires also tuning of the nonbonded terms.^10, 12, 19–21, 24, 28, 30, 32, 55^ Correcting errors in H-bonding and stacking interactions exclusively via formally intramolecular dihedral potentials is certainly a non-optimal approach. It has also been suggested that improvements of the existing pair-additive biomolecular *ffs* may profit from increase of flexibility of the parametrization, which can be achieved for example by breaking of the Lennard-Jones term combination rules (i.e., the NBfix approach) or by adding new simple terms to the *ff* form (the gHBfix approach, see Figure 1).^4, 12^

Unique primary solution NMR data are available for a set of RNA TNs and contemporary enhanced-sampling simulation methods allow to obtain well-converged ensembles of these model systems. Therefore, RNA TNs can be straightforwardly used to tune the RNA *ffs* by comparing conformational ensembles from simulations with NMR experimental data, although one should always keep in mind that there is a high risk of overfitting the *ff* towards RNA A-form. Recently, we have used the gHBfix approach to improve performance of the common χ_OL3CP_ AMBER RNA *ff* version by strengthening the –NH…N– base-base H-bonds and weakening the SPh H-bonds; this *ff* version is abbreviated as gHBfix19 in the present work.^12^

The gHBfix19 version visibly improved the RNA simulations without so far any detected side-effects.^12^ However, this *ff* version was still not fully satisfactory in reproducing the experimental data for RNA TNs. Here we further modify the gHBfix19 potential by tuning stability of H-bond interactions formed by the terminal nucleotides, namely the SPh and BPh interactions. The approach is abbreviated as tHBfix term. The idea to treat H-bonds formed by groups from terminal residues differently from H-bonds formed by equivalent groups of other nucleotides can be justified by several reasons. First, these groups have different parameters compared to the internal nucleotides in the parent RNA *ff*s. Second, the terminal nucleotides in reality may have somewhat different electronic-structure properties from the internal nucleotides. Finally, MD simulations of TNs are very significantly affected by interactions involving the terminal nucleotides while such interactions usually have insignificant effect on simulations of larger RNAs. Thus, considering the empirical nature of the *ff*, it is meaningful to not sacrifice transferability of the parameters of the internal nucleotides in order to perfect simulations of the short TNs.

We applied enhanced-sampling methods on the testing set containing five RNA TNs and compared the structural ensembles from simulations with accurate experimental datasets. The total raw amount of our simulations is over two milliseconds. Our results show that r(CAAU), r(CCCC) and r(GACC) TNs are mostly sampling A-form–like conformations in agreement, within the resolution limits, with the experiments. The remaining r(AAAA) and r(UUUU) TNs display a higher complexity with a mixture of structures. Comparison with the primary NMR data shows that their MD ensembles are likely different, to a detectable extent, from those populated in the experiment. However, it was not possible to unambiguously identify the exact origin of these residual discrepancies.

As expected based on the fact that we increased the number of parameters and that we used the NMR data in training them, the results with the newly designed χ_OL3CP_ + gHBfix19 + tHBfix20 potential are significantly improved in comparison with our previous work with gHBfix19.^12^ Although we did not make any extensive testing we suggest that side effects from adding the tHBfix20 parameters should be marginal since only terminal nucleotides are affected. Due to the profound sensitivity of TN simulations to the water models, the gHBfix19 + tHBfix20 parameter set is rather specific for the OPC water model. We also present set of NBfix parameters that should provide similar results as the gHBfix19 + tHBfix20 refinement.

In conclusion, we show that targeting the terminal residues of the *ff* may be considered both as an attempt to reflect potentially different electronic structures of these residues and a pragmatic means to increase tuning capability of the *ff*. Changing behavior of terminal nucleotides should not have side effects when simulating larger RNA motifs. Additionally, we propose that specific modifications of H-bonds formed by terminal nucleotides might potentially improve other *ff* features like spurious dynamics of RNA termini, description of base-pair fraying and simulations of protein-RNA complexes, where RNA self-interactions can create problems.^4^ In overall, the tHBfix potential is a generally transferable *ff* modification and can be applied to generic RNA motifs, although future refinements of the parameters are to be expected. This is because tuning of the non-bonded terms, paring them with water models and subsequent testing of all different *ff* versions on diverse RNAs represents a major computational challenge which cannot be fully accomplished within the framework of one computational study.

## Supporting information

Supporting Information for the article.

## ASSOCIATED CONTENT

### Supporting Information

The following files are available free of charge. Detailed description of MD protocol, Supporting Tables and Figures (PDF).

## ACKNOWLEDGMENT

Scott Kennedy and Douglas Turner are acknowledged for providing the NMR dataset for the r(CAAU) TN. This work was supported by the Operational Programme Research, Development and Education–European Regional Development Fund, Project No. CZ.02.1.01/0.0/0.0/16_019/0000754 (P.B., P.K., M.O.), Czech Science Foundation (18-25349S to P.B., P.K. and 20-16554S to V.M. and J.S.) and by Praemium Academiae (J.S.). This work was also supported by the project SYMBIT reg. number: CZ.02.1.01/0.0/0.0/15_003/0000477 financed by the ERDF (J.S.)

