## Supporting Information for the article. for "Fine-tuning of the AMBER RNA Force Field with a New Term Adjusting Interactions of Terminal Nucleotides"

**Additional description of the molecular dynamics protocol.** The RNA molecule remained constrained during minimization and optimization of waters and ions. Subsequently, all RNA atoms were frozen and the solvent molecules with counter-ions were allowed to move during a 500-ps long molecular dynamics (MD) run under NpT conditions (p = 1 atm., T = 298.16 K) in order to relax the total density. After this, the RNA molecule was relaxed by several minimization runs, with decreasing force constant applied to the sugar-phosphate backbone atoms. Subsequently, the system was heated in two steps: the first step involved heating under NVT conditions for 100 ps, whereas the second step involved density equilibration under NpT conditions for and additional 100 ps. The particle mesh Ewald (PME) method for treating electrostatic interactions was used. The standard unbiased MD simulations were performed under periodic boundary conditions in the NpT ensemble at 298.16 K using weak-coupling Berendsen thermostat[1] with coupling time of a 1 ps. The SHAKE algorithm, with a tolerance of $10^{-5}$ Å, was used to fix the positions of all hydrogen atoms, and a 10.0 Å cut-off was applied.

### Supporting Tables

**Table S1:** Overview of tested variants and combinations of the gHBfix and tHBfix potentials.[2] Each tested setting of the HBfix potentials is marked as $\mathrm{HBfix}\binom{\text{interaction pair}}{\eta}$, where the upper label in the parentheses defines the specific interactions between RNA groups and the lower parameter η defines the total energy modification for each H-bond interaction of this kind (in kcal/mol). [a]

| Label | gHBfix/tHBfix full notation [b] | Groups involved | |
|---|---|---|---|
| | | Donors | Acceptors |
| gHBfix19 | $\mathrm{gHBfix}\binom{2-\mathrm{OH}\ldots\mathrm{nbO/bO}}{-0.5}\binom{\mathrm{NH}\ldots\mathrm{N}}{+1.0}$ | 2'-OH/3'-OH/ 5'-OH (sugar), NH (base) | bO/nbO (phosphate), N (base) |
| gHBfix19$_{v2}$ [c] | $\mathrm{gHBfix}\binom{2-\mathrm{OH}\ldots\mathrm{nbO}}{-0.5}\binom{\mathrm{NH}\ldots\mathrm{N}}{+1.0}$ | 2'-OH/3'-OH/ 5'-OH (sugar), NH (base) | nbO (phosphate), N (base) |
| gHBfix19$_{v3}$ | $\mathrm{gHBfix}\binom{2-\mathrm{OH}\ldots\mathrm{nbO}}{-1.0}\binom{\mathrm{NH}\ldots\mathrm{N}}{+1.0}$ | 2'-OH/3'-OH/ 5'-OH (sugar), NH (base) | nbO (phosphate), N (base) |
| gHBfix19_NH-N$_{+0.75}$ | $\mathrm{gHBfix}\binom{2-\mathrm{OH}\ldots\mathrm{nbO/bO}}{-0.5}\binom{\mathrm{NH}\ldots\mathrm{N}}{+0.75}$ | 2'-OH/3'-OH/ 5'-OH (sugar), NH (base) | bO/nbO (phosphate), N (base) |
| gHBfix19$_{v2}$_BPh | $\mathrm{gHBfix}\binom{2-\mathrm{OH}\ldots\mathrm{nbO}}{-0.5}\binom{\mathrm{NH}\ldots\mathrm{N}}{+1.0}\binom{\mathrm{NH}\ldots\mathrm{nbO}}{-0.5}$ | 2'-OH/3'-OH/ 5'-OH (sugar), NH (base) | nbO (phosphate), N (base) |
| gHBfix19$_{v2}$_BPh$_{v2}$ | $\mathrm{gHBfix}\binom{2-\mathrm{OH}\ldots\mathrm{nbO}}{-0.5}\binom{\mathrm{NH}\ldots\mathrm{N}}{+1.0}\binom{\mathrm{NH}\ldots\mathrm{nbO}}{-1.0}$ | 2'-OH/3'-OH/ 5'-OH (sugar), NH (base) | nbO (phosphate), N (base) |
| gHBfix19$_{v3}$_BPh | $\mathrm{gHBfix}\binom{2-\mathrm{OH}\ldots\mathrm{nbO}}{-1.0}\binom{\mathrm{NH}\ldots\mathrm{N}}{+1.0}\binom{\mathrm{NH}\ldots\mathrm{nbO}}{-0.5}$ | 2'-OH/3'-OH/ 5'-OH (sugar), NH (base) | nbO (phosphate), N (base) |
| tHBfix(NH…nbO) [d] | $\mathrm{tHBfix}\binom{\mathrm{NH}\ldots\mathrm{nbO}}{-1.0}$ | NH (base) | nbO (phosphate) |
| tHBfix(OH…nbO) [d] | $\mathrm{tHBfix}\binom{2-\mathrm{OH}\ldots\mathrm{nbO}}{-1.0}$ | 2'-OH/3'-OH/ 5'-OH (sugar) | nbO (phosphate) |
| tHBfix(OH…O) [d] | $\mathrm{tHBfix}\binom{2-\mathrm{OH}\ldots\mathrm{O}}{-1.0}$ | 5'-OH (sugar) | O (base) |
| tHBfix(OH…OH) [d] | $\mathrm{tHBfix}\binom{2-\mathrm{OH}\ldots\mathrm{O2}}{-1.0}$ | 2'-OH/3'-OH (sugar) | 5'-OH (sugar) |
| tHBfix20 [e] | $\mathrm{tHBfix}\binom{2-\mathrm{OH}\ldots\mathrm{nbO/O/O2}}{-1.0}\binom{\mathrm{NH}\ldots\mathrm{nbO}}{-1.0}$ | 2'-OH/3'-OH/ 5'-OH (sugar), NH (base) | nbO (phosphate), N/O (base), 5'-OH (sugar) |

[a] see the original paper[2] for the detailed definition of the gHBfix potential and for the complete list of atoms from each RNA nucleotide whose interactions are modified by the gHBfix. All listed potentials are applied between hydrogen of the proton donor and proton acceptor heavy atom. In the present study we applied the HBfix terms in all cases using the interval between 2 Å and 3 Å, see Figure 1 in the main text.

[b] as used in the original paper introducing the gHBfix potential.[2]

[c] This variant is marginally different from the basic gHBfix19 version by not modulating interactions between the 2'-OH groups and bridging phosphate oxygens (bO). We assume the difference between gHBfix19 and gHBfix19$_{v2}$ versions has minimal impact on the simulations, due to the marginal role of the interactions involving the bridging oxygens.

[d] see Table 1 in the main text for the detailed list of atoms from RNA nucleotides whose interactions were modified.

[e] containing the combination of all above-introduced tHBfix terms. See the Table S2 with enumeration of all the interactions.

**Table S2:** List of interactions that were modified by the tHBfix20 potential for the particular TN motif.

| TN | H-bonds |
|---|---|
| GACC | $C_{S4}$(2'-OH/3'-OH)…$C_{S3}$(*pro*-$R_P$/*pro*-$S_P$) |
| | $G_{S1}$(5'-OH)…$C_{S3}$(*pro*-$R_P$/*pro*-$S_P$) |
| | $G_{S1}$(5'-OH)…$C_{S4}$(O2) |
| | $C_{S4}$(2'-OH/3'-OH)…$G_{S1}$(O5') |
| CAAU | $C_1$(N4H)…$U_{S4}$(*pro*-$R_P$/*pro*-$S_P$) |
| | $U_{S4}$(2'-OH/3'-OH)…$A_{S3}$(*pro*-$R_P$/*pro*-$S_P$) |
| | $C_{S1}$(5'-OH)…$A_{S3}$(*pro*-$R_P$/*pro*-$S_P$) |
| | $C_{S1}$(5'-OH)…$U_{S4}$(O2) |
| | $U_{S4}$(2'-OH/3'-OH)…$C_{S1}$(O5') |
| AAAA | $A_1$(N6H)…$A_{S4}$(*pro*-$R_P$/*pro*-$S_P$) |
| | $A_{S4}$(2'-OH/3'-OH)…$A_{S3}$(*pro*-$R_P$/*pro*-$S_P$) |
| | $A_{S1}$(5'-OH)…$A_{S3}$(*pro*-$R_P$/*pro*-$S_P$) |
| | $A_{S4}$(2'-OH/3'-OH)…$A_{S1}$(O5') |
| CCCC | $C_1$(N4H)…$C_{S4}$(*pro*-$R_P$/*pro*-$S_P$) |
| | $C_{S4}$(2'-OH/3'-OH)…$C_{S3}$(*pro*-$R_P$/*pro*-$S_P$) |
| | $C_{S1}$(5'-OH)…$C_{S3}$(*pro*-$R_P$/*pro*-$S_P$) |
| | $C_{S1}$(5'-OH)…$C_{S4}$(O2) |
| | $C_{S4}$(2'-OH/3'-OH)…$C_{S1}$(O5') |
| UUUU | $U_{S4}$(2'-OH/3'-OH)…$U_{S3}$(*pro*-$R_P$/*pro*-$S_P$) |
| | $U_{S1}$(5'-OH)…$U_{S3}$(*pro*-$R_P$/*pro*-$S_P$) |
| | $U_{S1}$(5'-OH)…$U_{S4}$(O2) |
| | $U_{S4}$(2'-OH/3'-OH)…$U_{S1}$(O5') |

**Table S3:** The stacking overlap (in relative values) between consecutive nucleobases from the simulation of the r(AAAA) TN and its dependency on the *syn*/*anti* combination of nucleobases (see Methods in the main text for details).

| $\chi$-dihedral major states | 5'-end ($A_{S1}$-$A_{S2}$) | middle ($A_{S2}$-$A_{S3}$) | 3'-end ($A_{S3}$-$A_{S4}$) |
|---|---|---|---|
| *anti*/*anti* | 0.31 | 0.30 | 0.30 |
| *anti*/*syn* | 0.21 | 0.25 | 0.25 |
| *syn*/*anti* | 0.20 | 0.30 | 0.41 |
| *syn*/*syn* | 0.14 | 0.38 | 0.29 |

**Table S4:** The modified Lennard-Jones combining rules (NBfix) obtained as an attempt to mimic modulation of the interaction-energy curves introduced by the tHBfix20 potential. The NBfix parameters mimicking modulation of the gHBfix19 potential (included thus also in the combined gHBfix19 + tHBfix20 version) and other details can be found in the Supporting Information of Ref. [2].

| tHBfix20 [a] | η [b] (kcal/mol) | Atom pairs modified [c] | $R$ (Å) [d] $\chi_{OL3CP}$ [f] | NBfix mimicking tHBfix20 [g] | $\varepsilon$ (kcal/mol) [e] $\chi_{OL3CP}$ [f] | NBfix mimicking tHBfix20 [g] |
|---|---|---|---|---|---|---|
| NH…nbO | –1.0 | HN(H)…OX(OP) | 2.3493 | 2.4993 | 0.0574 | 0.1449 |
| 2-OH…bO | –1.0 | HT(HO)…OT(OR) | 1.7718 [h] | 2.3918 | 0 | 0.17 |
| 2-OH…nbO | –1.0 | HT(HO)…OX(OP) | 1.7493 [h] | 2.1693 | 0 | 0.1889 |
| 2-OH…O | –1.0 | HT(HO)…OU(O) | 1.6612 [h] | 2.1912 | 0 | 0.1889 |

[a] interactions modified by the tHBfix20 potential

[b] total potential energy modulation for each H-bond interaction by the tHBfix potential function (see Methods in the main text for details and Figure 1 in the main text for definition of the η parameter; its negative value means that the interaction is penalized).

[c] new atom types were designed in order to differentiate among interactions formed by groups from terminal and internal nucleotides (original atom types from the $\chi_{OL3CP}$ *ff* are in parenthesis)

[d] distance, where the Lennard-Jones potential for the interaction of atoms $i$ and $j$, $R_{i,j}$, is exactly zero. Parameters the particular Lennard-Jones pairs are derived as: $R_{i,j} = R_i + R_j$

[e] depth of the potential well for the interaction of atoms $i$ and $j$, $\varepsilon_{i,j}$. Parameters for the particular Lennard-Jones pairs are derived as: $\varepsilon_{i,j} = \sqrt{(\epsilon_i + \varepsilon_j)}$

[f] *ff*99bsc0$\chi_{OL3}$[3-6] RNA *ff* version with the vdW modification of phosphate oxygens developed by Steinbrecher et al.[7]

[g] NBfix *ff* reparameterization was prepared in a way to be comparable with the tHBfix$\binom{2\text{–OH…nbO/O/O2}}{-1.0}\binom{\text{NH…nbO}}{-1.0}$ potential (i.e. tHBfix20, see Table S1).

[h] in the original *ff* these values are formally equal to radii of the bO, nbO and carbonyl group oxygens, respectively, as the radius of polar hydrogen is equal to zero. However, they are essentially irrelevant due to zero $\varepsilon$ value.

**Supporting Figures**

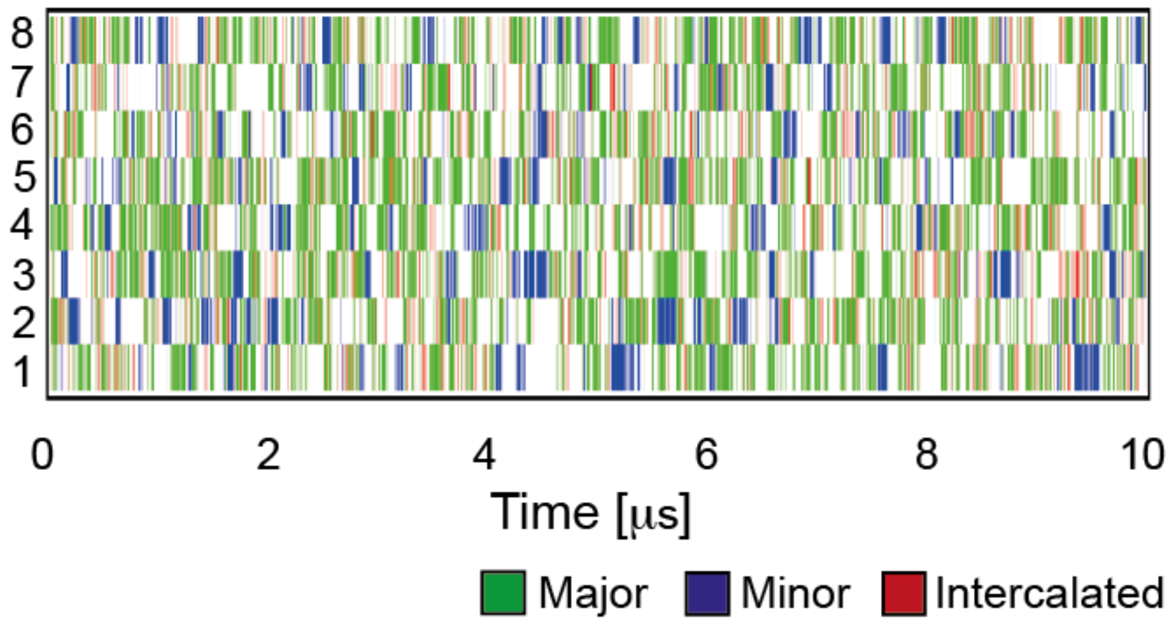

**Figure S1:** A representative example of conformational sampling and convergence of one REST2 simulation, namely the REST2 simulation of the r(CCCC) TN with the combined gHBfix19 + tHBfix(NH…nbO) potential. Time evolution of major conformers, i.e., A-RNA major (green), A-RNA minor (blue), and the spurious intercalated structure (red) in all eight replicas analyzed from continuous (demultiplexed) trajectories.

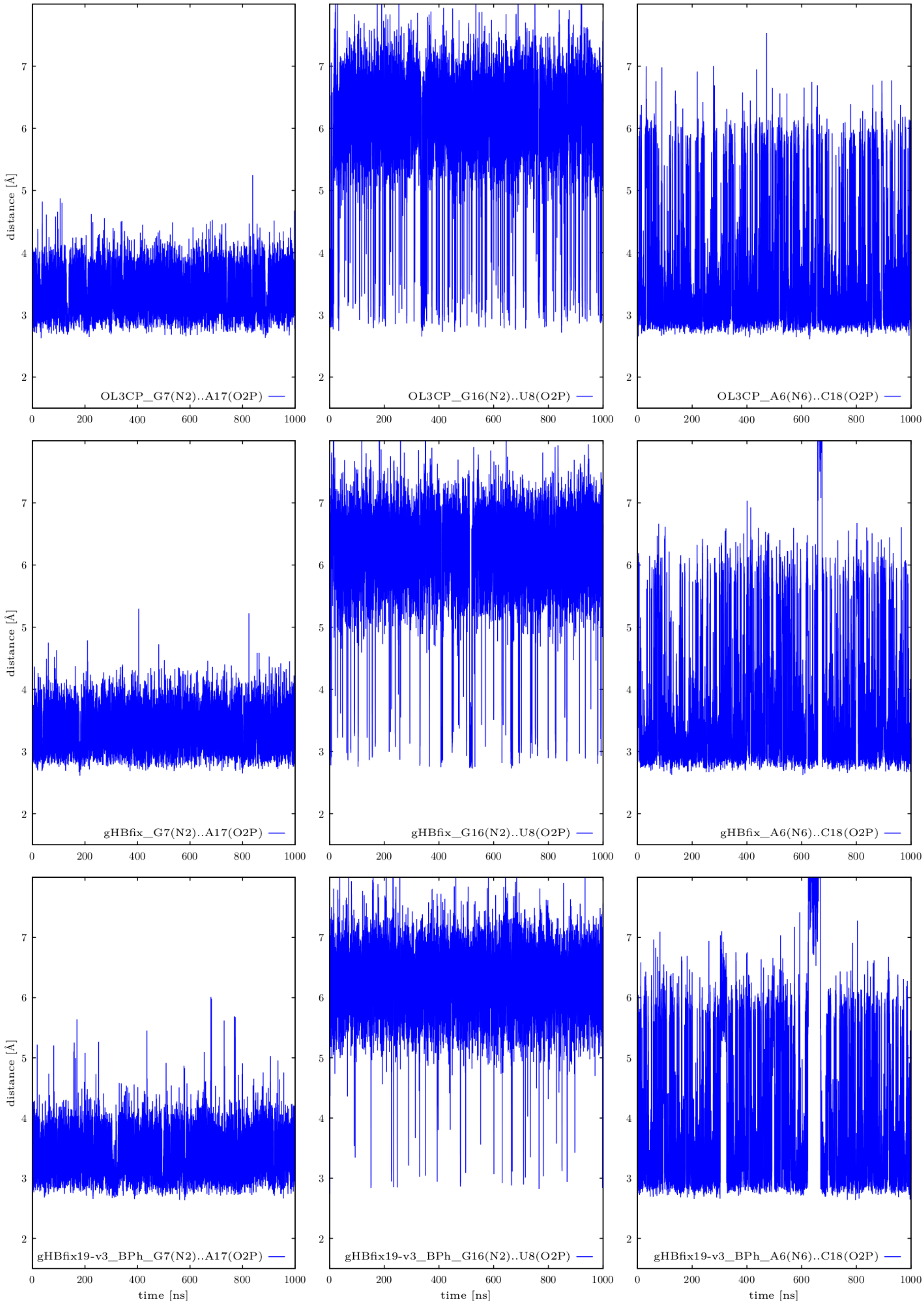

**Figure S2:** Structural analysis of three standard MD simulations of Sarcin-Ricin RNA loop (SRL) motif. See Figure 1 in Ref. [8] for the structure. Plots in the first, second, and third column display fluctuations of distances indicating signature BPh interactions, i.e., G7(N2)…A17(*pro*-$R_P$), G16(N2)…U8(*pro*-$R_P$), and A6(N6)…C18(*pro*-$R_P$) distances, respectively. We compared the behavior in three different RNA *ff*s during 1 μs long MD simulations, i.e., the standard $\chi_{OL3CP}$[3-6] (OL3CP, plots at the top), $\chi_{OL3CP}$ with the gHBfix19 potential[2] (gHBfix, plots in the middle), and $\chi_{OL3CP}$ with the gHBfix19 combined with –0.5 kcal/mol penalty of all possible BPh interactions and with increased weakening of all the SPh interactions to –1.0 kcal/mol (gHBfix19$_{v3}$_BPh, plots at the bottom).

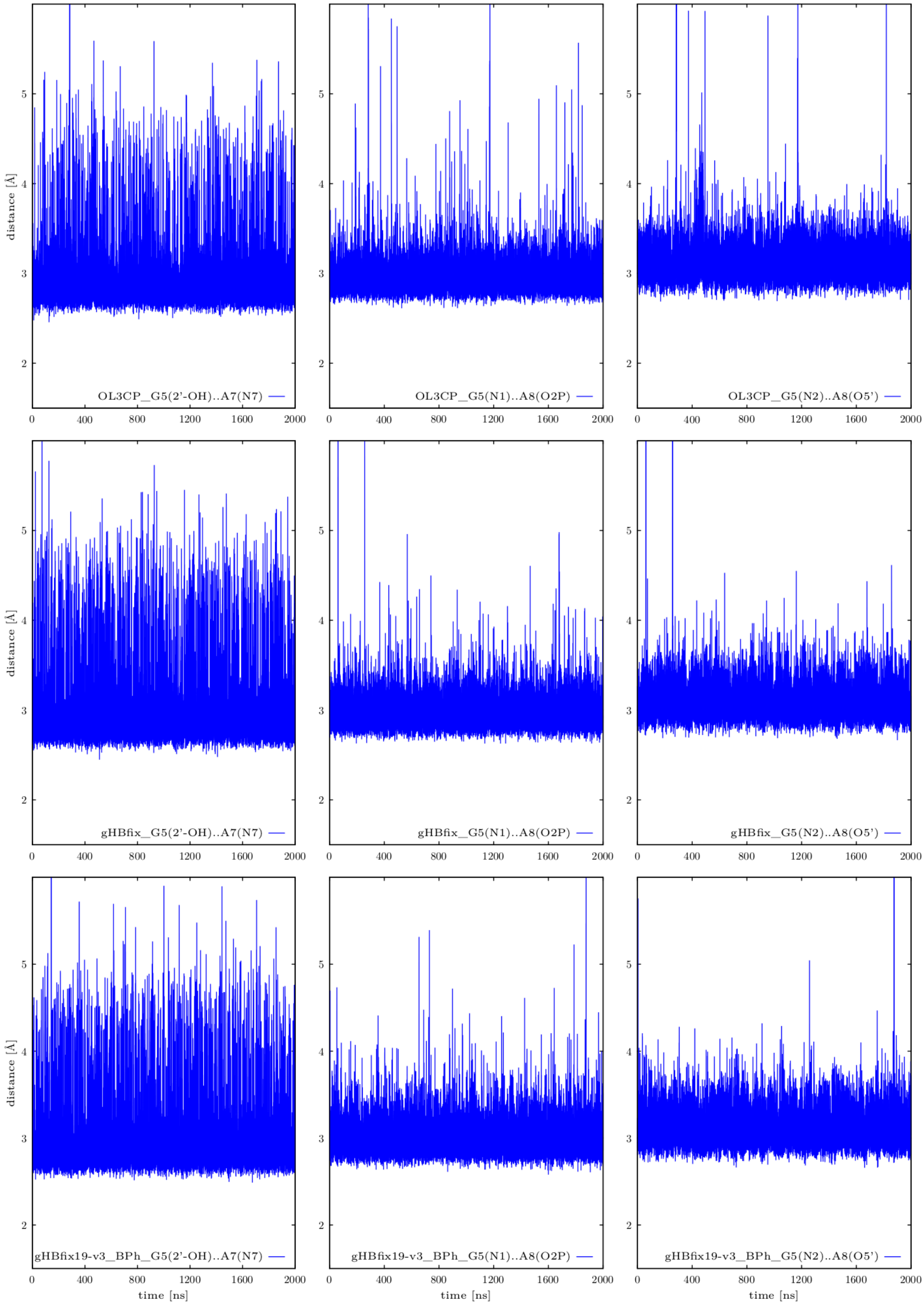

**Figure S3:** Structural analysis of three unbiased MD simulations of T-loop RNA motif. See Figure 1 in Ref. [9] for the structure. Plots in the first, second, and third column display fluctuations of distances indicating signature H-bonds, i.e., G5(2'-OH)…A7(N7), G5(N1)…A8(*pro*-$R_P$), and G5(N2)…A8(O5') distances, respectively. We compared the behavior in three different RNA *ff*s during 2 μs long MD simulations, i.e., the standard $\chi_{OL3CP}$[3-6] (OL3CP, plots at the top), $\chi_{OL3CP}$ with the external gHBfix19 potential[2] (gHBfix, plots in the middle), and $\chi_{OL3CP}$ with the gHBfix19 combined with –0.5 kcal/mol penalty of all possible BPh interactions and with increased weakening of all the SPh interactions to –1.0 kcal/mol (gHBfix19$_{v3}$_BPh, plots at the bottom).

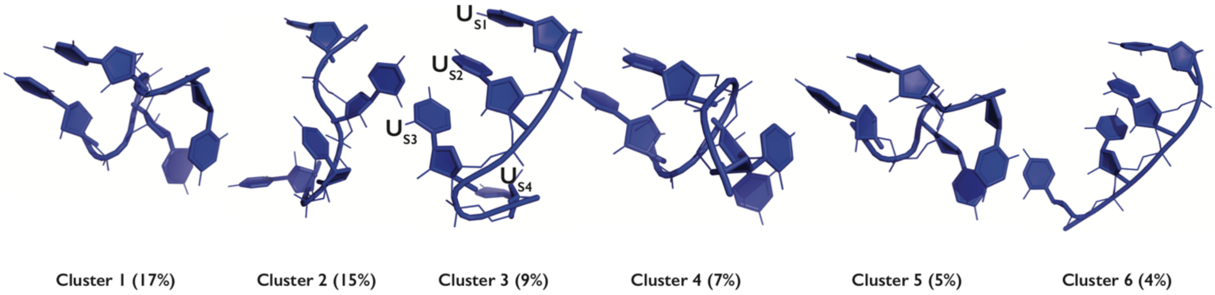

**Figure S4:** Tertiary structures of the most populated clusters from r(UUUU) REST2 simulation with the $\chi_{OL3CP}$ RNA *ff* modified by the combination of gHBfix19[2] + tHBfix20 (present work, Table S1) external potentials. H-atoms are not shown for clarity. Population of unassigned structures is ~32%.

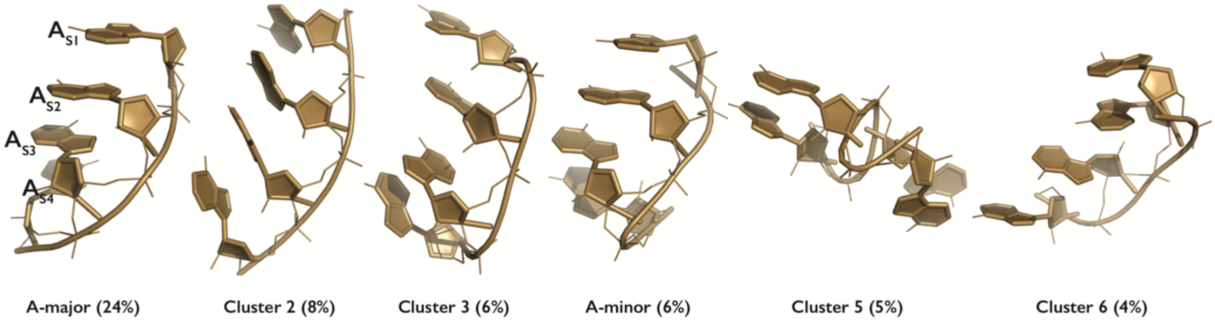

**Figure S5:** Tertiary structures of the most populated clusters from r(AAAA) REST2 simulation with the $\chi_{OL3CP}$ RNA *ff* modified by the combination of gHBfix19[2] + tHBfix20 (Table S1) potentials. H-atoms are not shown for clarity.

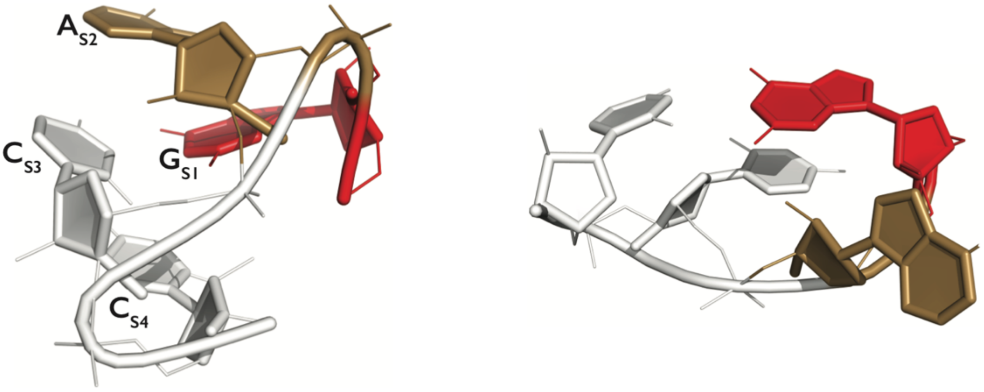

1-3 loop 1-4 loop

**Figure S6:** Tertiary structures of the loop-like clusters of the r(GACC) TN, which are commonly occurring during REST2 simulation (especially with three-point charge water models). A, C, and G nucleotides are colored in sand, white, and red, respectively. H-atoms are not shown for clarity. The loop-like clusters indicate overestimation of base pairing.
